# Predicting long-term multicategory cause of death in patients with prostate cancer: random forest versus multinomial model

**DOI:** 10.1101/2020.01.03.893966

**Authors:** Jianwei Wang, Fei Deng, Fuqing Zeng, Andrew J. Shanahan, Wei V. Li, Lanjing Zhang

## Abstract

Patients with prostate cancer more likely die of non-cancer cause of death (COD) than prostate cancer. It is thus important to accurately predict multi-category COD in these patients. Random forest (RF), a popular machine learning model, has been shown useful for predicting binary cancer-specific deaths. However, its accuracy for predicting multi-category COD in cancer patients is unclear. We included patients in Surveillance, Epidemiology, and End Results-18 cancer registry-program with prostate cancer diagnosed in 2004 (followed-up through 2016). They were randomly divided into training and testing sets with equal sizes. We evaluated prediction accuracies of RF and conventional-statistical/multinomial models for 6-category COD by data-encoding types using the 2-fold cross-validation approach. Among 49,864 prostate cancer patients, 29,611 (59.4%) were alive at the end of follow-up, and 5,448 (10.9%) died of cardiovascular disease, 4,607 (9.2%) of prostate cancer, 3,681 (7.4%) of Non-Prostate cancer, 717 (1.4%) of infection, and 5,800 (11.6%) of other causes. We predicted 6-category COD among these patients with a mean accuracy of 59.1% (n=240, 95% CI, 58.7%-59.4%) in RF models with one-hot encoding, and 50.4% (95% CI, 49.7%-51.0%) in multinomial models. Tumor characteristics, prostate-specific antigen level, and diagnosis confirmation-method were important in RF and multinomial models. In RF models, no statistical differences were found between the accuracies of development versus cross validation phases, and those of categorical versus one-hot encoding. We here report a RF model that has an accuracy of 59.1% in predicting long-term 6-category COD among prostate cancer patients. It outperforms multinomial logistic models (absolute prediction-accuracy difference, 8.7%).

## Introduction

Prostate cancer is the most prevalent cancer and the second leading-cause of cancer deaths among men in the U.S.A., accounting for 174,650 new cases and 31,620 deaths in 2019.[1, 2] More patients with prostate cancer died of non-cancer causes than of prostate cancer.[3, 4] It is thus important to understand, predict and prevent non-cancer causes of death (CODs) among these patients, particularly cardiovascular disease (CVD).[5] However, only a limited number of studies investigated multi-category COD in prostate cancer patients, and none of them were focused on the prediction of COD.[3, 5, 6]

The random forest (RF) model, a widely-used machine/statistical learning model, improves the performance of decision trees through random sampling of training data when building trees and random subsetting of features when splitting nodes.[7] The RF model often outperforms several machine learning and conventional statistical (e.g. logistic regression) models in predicting binary cancer-specific or all-cause deaths, [8-12] with exceptions in a few simulation or biomarker studies.[13, 14] It has also been used to predict cancer-specific deaths in prostate cancer patients.[15] However, few studies have used RF model for predicting multi-category COD in cancer patients, or compared the prediction accuracies of RF versus conventional statistical model (e.g. multinomial logistic regression) for multi-category COD. Our research aims to fill this gap, and we designed a population-based observational study to predict 12-year multi-category COD in prostate cancer patients using RF and multinomial logistic models.

## Methods

### Patient Data

We extracted individual-patient data from the Surveillance, Epidemiology, and End Results-18 (SEER-18) Program (www.seer.cancer.gov) SEER*Stat Database with Treatment Data using the SEER*Stat software (Surveillance Research Program, National Cancer Institute SEER*Stat software (seer.cancer.gov/seerstat) version <8.3.6>).[16] SEER-18 is the largest SEER database including cases from 18 states and covers near 30% of the U.S. population.[17] The datasets have been widely used and validated for research on breast and colorectal cancers.[18-20] Any summary data involving fewer than 15 patients were statistically suppressed to protect patient identity. Since the SEER database is an existing, de-identified and publicly available dataset, this study is exempt from Institutional Review Board (IRB) review under exempt category 4.

We included all qualified invasive prostate cancer cases in SEER-18 diagnosed in 2004 (2019 data-release, followed up through December 2016). The diagnosis year of 2004 was chosen because the 6 ^th^ edition of the Tumor, Node and Metastasis staging manual (TNM6) of the American Joint Commission on Cancer (AJCC) was started in 2004 and allowed 12 years of follow-up. But, the AJCC 7 ^th^ edition of the Tumor, Node and Metastasis staging manual (TNM7) was started in 2010, and would allow only up to 6 years of follow-up, which was not long enough in our view. The inclusion criteria were survival time longer than 1 month, aged 20 years and older, with known COD and first primary only.

### Outcome and Variables

The outcome of the statistical models was the patients’ 6-category COD. The COD were originally classified using SEER’s recodes of the causes of death according to the COD definition of the U.S. Mortality Data, which were extracted from underlying cause of death on the death certificates of deceased patients.[21] The underlying COD was the unique and most important etiology of the patients’ death, while other causes may link to the death and be recorded as other COD on the death certificate. We simplified the SEER COD into 6 categories based on the prevalence of COD,[3, 6, 15] including alive, CVD, infection, non-prostate cancer, prostate cancer and others.

The following factors were included in the analysis as variables in RF or multinomial models: age at diagnosis, race/ethnicity (non-Hispanic White, Hispanic, non-Hispanic Black, Asian and Pacific Islanders, and others),[22] T, N and M categories of TNM6, AJCC TNM6 clinical staging, prostate specific antigen level (PSA, ng/ml), sum of the Gleason score, chemotherapy, radiotherapy, surgery, and attributes of the county where the patient resided at the time of diagnosis.[23] The PSA levels and Gleason scores were collected from medical records as site specific factors of prostate cancer since 2010.[24, 25] Specifically, sums of the Gleason score were obtained from pathology report of resected specimen when available, or that of biopsy specimen if no surgery done. The 4 census-regions of patient’s residence county were defined by the U.S. Census Bureau.[26] We converted continuous variables into 4-category variables based on their quartiles. The chemotherapy and radiotherapy data were obtained after signing a user agreement.[25, 27] It is noteworthy that no or unknown status of these treatments should be considered less reliable, while receipt of these treatments was generally confirmed and reliable. [25, 27]

### Statistical analysis

We compared the accuracies of the RF and multinomial logit models after tuning the parameters of the RF model and choosing the model with the best accuracy. Using the two-fold cross-validation approach, the patients were first randomly divided into two subsets of similar sizes (n=25,000 and 24,864, respectively). In each round of the validation, one subset is treated as the training data for constructing models, and the other subset is treated as the test data for evaluating prediction performance (**Figure 1**). For data-quality assurance, we compared the covariates in the training and testing sets using Chi-square or Student’s t test. We identified the RF model with the best accuracy, which is termed as tuning process in data science. Specifically, we examined prediction accuracies (i.e. 1 – classification error) of the models with various numbers of iterations (from 50 to 800 by an interval of 50) and variables (from 1 to 15), which were the number of computation rounds and the preset number of the features in RF model, respectively. After the two rounds of validation, the set of parameters that led to the RF model with the smallest average classification error were selected. This cross-validation process is outlined in **Figure 1**.

**Figure 1.**
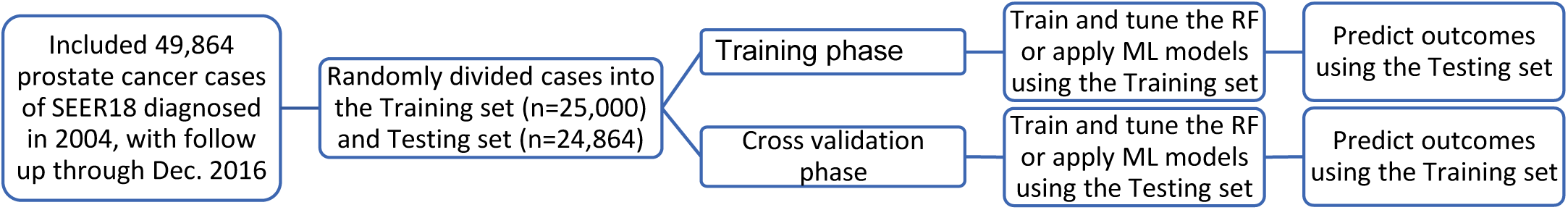
Study flow. We randomized the patients into training and testing sets with similar sample-size in each group.We then tuned the random forest (RF) model, chose the best-fit RF model, and applied multinomial logistic (ML) model using training set. Using the ML and chosen RF models, we predicted 6-category causes of death among the patients in testing set. During the cross validation phase, we followed a similar protocol but with swapped datasets.

Several sensitivity analyses were performed on RF models. To exclude patients lost to follow-up, we conducted training and validation processes in the patients who died during the follow-up or was alive for >150 months (12.5 years). We also generated training and testing sets with balanced distribution in all tested independent and dependent variables, which were assessed using Chi-square or Student’s t test by each variable.

A previous study has shown that one-hot encoding could sometimes outperform complex encoding systems.[28] This approach was also used in machine learning models of cancer driver genes.[29] For one-hot encoding, all multicategory variables (i.e. discrete variables with more than two catrgories) were transformed into a new set of binary variables. For example, the categorical variable for race/ethnicity group would be replaced by 5 binary variables representing whether the patients are non-Hispanic White, Hispanic, non-Hispanic Black, Asian and Pacific Islanders, or others, respectively. We trained the RF or multinomial logit models using the one-hot encoded data, and compared the results with those using multicategory variables.

For the multinomial logistic regression model, we first constructed the model using the training set (the subset with sample size of 25,000) and predicted the 6-category COD using testing set (**Figure 1**). If the predicted probability of a given COD was higher than 0.5, the COD would be assigned to the COD of the patient. Ideally, only one COD had a predicted probability >0.5 and was allowed for each case, thus any patient with 0 or >1 predicted COD was considered unsuccessfully predicted using multinomial model.

We carried out the above statistical analyses using the RF package and multinomial logistic models of Stata (version 16, College Station, TX).[30-32] The 95% confidence intervals (CI) of prediction accuracies were estimated using both binomial and Poisson models, that produced very similar results. All P values were two-tailed, and a P value <0.05 was considered statistically significant.

## Results

### Patients

We identified and analyzed 49,864 men with prostate cancer diagnosed in 2004 in the SEER-18 (**Table 1**), including 29,611 (59.4%) alive, 5,448 (10.9) died of CVD, 4,607 (9.2%) of prostate cancer, 3,681 (7.4%) of non-prostate cancer, 717 (1.4%) of infection, and 5,800 (11.6%) of other causes. The mean survival time was 117 months, while there were 31,273 patients who died during followup or was alive for >150 months. Majority of the cancers were of AJCC 6 stage 2 (80.9%) and not treated with prostatectomy (61.6%). We randomly divided the cases into training and testing sets (**STable 1**), and found the outcome and all covariates were similarly distributed in these sets, except radiotherapy status (P=0.047). We then sorted the data by outcome and radiotherapy, randomized the cases again, and achieved similar distributions of the outcome and on covariates in the two sets (**STable 2**). For the sensitivity analyses on the patients who died during followup or was alive >150 months, CODs were similarly distributed in the training and testing sets (**Stable 3**).

**Table 1.** Baseline characteristics of included subjects.

|  | Alive,<br>n=29,611 | CVD,<br>n=5,448 | Infection,<br>n=717 | Non-Prostate<br>cancer,<br>n=3,681 | Other<br>cause,<br>n=5,800 | Prostate<br>cancer,<br>n=4,607 | Total,<br>n=49,864 |
| --- | --- | --- | --- | --- | --- | --- | --- |
| **Age (yr)¶** | 63 (50-77) | 74 (59-87) | 75 (58-88) | 70 (56-83) | 73 (58-86) | 72 (54-88) | 67 (51-83) |
| **Survival time (mo)¶** | 146 (131-155) | 77 (7-143) | 78 (6-141) | 78 (12-141) | 82 (11-143) | 59 (4-137) | 117 (16-154) |
| **Race** |  |  |  |  |  |  |  |
| API | 1,453 | 210 | 43 | 172 | 268 | 195 | 2,341 |
| (%) | (4.9) | (3.9) | (6.0) | (4.7) | (4.6) | (4.2) | (4.7) |
| Hispanic | 2,662 | 412 | 68 | 249 | 497 | 423 | 4,311 |
| (%) | (9.0) | (7.6) | (9.5) | (6.8) | (8.6) | (9.2) | (8.7) |
| NH Black | 3,830 | 865 | 143 | 553 | 807 | 812 | 7,010 |
| (%) | (12.9) | (15.9) | (19.9) | (15.0) | (13.9) | (17.6) | (14.1) |
| NH White | 21,093 | 3,920 | 461 | 2,690 | 4,189 | 3,147 | 35,500 |
| (%) | (71.2) | (72.0) | (64.3) | (73.1) | (72.2) | (68.3) | (71.2) |
| Unknown/Other | 573 | 41 | <15* | 17 | 39 | 30 | 702 |
| (%) | (1.9) | (0.8) |  | (0.5) | (0.7) | (0.7) | (1.4) |
| **TNM6 T category** |  |  |  |  |  |  |  |
| T1/2 | 26,641 | 4,873 | 635 | 3,245 | 5,201 | 2,917 | 43,512 |
| (%) | (90.0) | (89.5) | (88.6) | (88.2) | (89.7) | (63.3) | (87.3) |
| T3/4 | 2,543 | 278 | 40 | 281 | 329 | 890 | 4,361 |
| (%) | (8.6) | (5.1) | (5.6) | (7.6) | (5.7) | (19.3) | (8.8) |
| Unknown/Other | 427 | 297 | 42 | 155 | 270 | 800 | 1,991 |
| (%) | (1.4) | (5.5) | (5.9) | (4.2) | (4.7) | (17.4) | (4.0) |
| **TNM6 N category** |  |  |  |  |  |  |  |
| 0 | 28,140 | 4,850 | 631 | 3,354 | 5,226 | 3,057 | 45,258 |
| (%) | (94.7) | (88.3) | (87.2) | (90.6) | (89.4) | (65.2) | (90.2) |
| 1 | 283 | 64 | <15* | 60 | 55 | 357 | 830 |
| (%) | (1.0) | (1.2) |  | (1.6) | (0.9) | (7.6) | (1.7) |
| Unknown/Other | 1,307 | 579 | 82 | 289 | 566 | 1,272 | 4,095 |
| (%) | (4.4) | (10.5) | (11.3) | (7.8) | (9.7) | (27.1) | (8.2) |
| **TNM6 M category** |  |  |  |  |  |  |  |
| 0 | 28,615 | 4,911 | 648 | 3,389 | 5,291 | 2,794 | 45,648 |
| (%) | (96.3) | (89.4) | (89.5) | (91.5) | (90.5) | (59.6) | (91.0) |
| 1 | 160 | 182 | 25 | 120 | 174 | 1,363 | 2,024 |
| (%) | (0.5) | (3.3) | (3.5) | (3.2) | (3.0) | (29.1) | (4.0) |
| Unknown/Other | 955 | 400 | 51 | 194 | 382 | 529 | 2,511 |
| (%) | (3.2) | (7.3) | (7.0) | (5.2) | (6.5) | (11.3) | (5.0) |
| **AJCC6 staging** |  |  |  |  |  |  |  |
| 1 | 47 | 21 | <15* | <15* | 26 | <15* | 110 |
| (%) | (0.2) | (0.4) |  |  | (0.5) |  | (0.2) |
| 2 | 25476 | 4459 | 574 | 3013 | 4785 | 2054 | 40361 |
| (%) | (86.0) | (81.9) | (80.1) | (81.9) | (82.5) | (44.6) | (80.9) |
| 3 | 2110 | 173 | 19 | 200 | 216 | 328 | 3046 |
| (%) | (7.1) | (3.2) | (2.7) | (5.4) | (3.7) | (7.1) | (6.1) |
| 4 | 607 | 252 | 38 | 200 | 240 | 1595 | 2932 |
| (%) | (2.1) | (4.6) | (5.3) | (5.4) | (4.1) | (34.6) | (5.9) |
| Unknown/Other | 1371 | 543 | 81 | 258 | 533 | 629 | 3415 |
| (%) | (4.6) | (10.0) | (11.3) | (7.0) | (9.2) | (13.7) | (6.9) |
| **Chemotherapy** |  |  |  |  |  |  |  |
| None/Unknown | 29,617 | 5,472 | 720 | 3,671 | 5,821 | 4,516 | 49,817 |
| (%) | (99.6) | (99.6) | (99.5) | (99.1) | (99.6) | (96.4) | (99.3) |
| Received | 113 | 21 | <15* | 32 | 26 | 170 | 366 |
| (%) | (0.4) | (0.4) |  | (0.9) | (0.4) | (3.6) | (0.7) |
| **Radiotherapy** |  |  |  |  |  |  |  |
| None/Unknown | 18,450 | 3,364 | 446 | 2,094 | 3,537 | 3,257 | 31,148 |
| (%) | (62.1) | (61.2) | (61.6) | (56.6) | (60.5) | (69.5) | (62.1) |
| Received | 11,280 | 2,129 | 278 | 1,609 | 2,310 | 1,429 | 19,035 |
| (%) | (37.9) | (38.8) | (38.4) | (43.5) | (39.5) | (30.5) | (37.9) |
| **Surgery** |  |  |  |  |  |  |  |
| Local Excision | 1,093 | 599 | 72 | 270 | 657 | 483 | 3,174 |
| (%) | (3.7) | (11.0) | (10.0) | (7.3) | (11.3) | (10.5) | (6.4) |
| No surgery | 15,142 | 4,261 | 578 | 2,649 | 4,413 | 3,666 | 30,709 |
| (%) | (51.1) | (78.2) | (80.6) | (72.0) | (76.1) | (79.6) | (61.6) |
| Prostatectomy | 13,376 | 588 | 67 | 762 | 730 | 458 | 15,981 |
| (%) | (45.2) | (10.8) | (9.3) | (20.7) | (12.6) | (9.9) | (32.1) |
| **Rural-urban continuum 2003\$** | | | | | | | |
| Metro | 26,709 | 4,758 | 635 | 3,178 | 4,958 | 4,039 | 44,277 |
| (%) | (89.8) | (86.6) | (87.7) | (85.8) | (84.8) | (86.2) | (88.2) |
| Non-Metro | 3,021 | 735 | 89 | 525 | 889 | 647 | 5,906 |
| (%) | (10.2) | (13.4) | (12.3) | (14.2) | (15.2) | (13.8) | (11.8) |
| **Census region** |  |  |  |  |  |  |  |
| Midwest | 2,946 | 658 | 77 | 399 | 613 | 429 | 5,122 |
| (%) | (10.0) | (12.1) | (10.7) | (10.8) | (10.6) | (9.3) | (10.3) |
| Northeast | 4,797 | 882 | 123 | 631 | 874 | 721 | 8,028 |
| (%) | (16.2) | (16.2) | (17.2) | (17.1) | (15.1) | (15.7) | (16.1) |
| South | 5,573 | 1,140 | 176 | 843 | 1,393 | 1,009 | 10,134 |
| (%) | (18.8) | (20.9) | (24.6) | (22.9) | (24.0) | (21.9) | (20.3) |
| West | 16,295 | 2,768 | 341 | 1,808 | 2,920 | 2,448 | 26,580 |
| (%) | (55.0) | (50.8) | (47.6) | (49.1) | (50.3) | (53.1) | (53.3) |
| **Percent of education attainment, quartile\$** | | | | | | | |
| Q1, <15.08 | 8,001 | 1,200 | 140 | 836 | 1,339 | 1,029 | 12,545 |
| (%) | (26.9) | (21.9) | (19.3) | (22.6) | (22.9) | (22.0) | (25.0) |
| Q2, 15.09-18.15 | 7,538 | 1,287 | 182 | 898 | 1,448 | 1,193 | 12,546 |
| (%) | (25.4) | (23.4) | (25.1) | (24.3) | (24.8) | (25.5) | (25.0) |
| Q3, 18.17-25.79 | 7,236 | 1,420 | 189 | 997 | 1,492 | 1,212 | 12,546 |
| (%) | (24.3) | (25.9) | (26.1) | (26.9) | (25.5) | (25.9) | (25.0) |
| Q4, >50.77 | 6,955 | 1,586 | 213 | 972 | 1,568 | 1,252 | 12,546 |
| (%) | (23.4) | (28.9) | (29.4) | (26.3) | (26.8) | (26.7) | (25.0) |
| **Percent of persons in poverty, quartile\$** | | | | | | | |
| Q1, <21.18 | 8,034 | 1,210 | 160 | 865 | 1,305 | 1,044 | 12,618 |
| (%) | (27.0) | (22.0) | (22.1) | (23.4) | (22.3) | (22.3) | (25.1) |
| Q2, 21.33-29.81 | 7,655 | 1,258 | 152 | 929 | 1,364 | 1,129 | 12,487 |
| (%) | (25.8) | (22.9) | (21.0) | (25.1) | (23.3) | (24.1) | (24.9) |
| Q3, 29.86-37.36 | 7,276 | 1,493 | 220 | 986 | 1,662 | 1,256 | 12,893 |
| (%) | (24.5) | (27.2) | (30.4) | (26.6) | (28.4) | (26.8) | (25.7) |
| Q4, >67.40 | 6,765 | 1,532 | 192 | 923 | 1,516 | 1,257 | 12,185 |
| (%) | (22.8) | (27.9) | (26.5) | (24.9) | (25.9) | (26.8) | (24.3) |
| **Percent of foreign-born residents, quartile§** |  |  |  |  |  |  |  |
| Q1, <5.95 | 6,864 | 1,467 | 188 | 1,041 | 1,747 | 1,254 | 12,561 |
| (%) | (23.1) | (26.7) | (26.0) | (28.1) | (29.9) | (26.8) | (25.0) |
| Q2, 5.98-15.22 | 7,739 | 1,399 | 172 | 922 | 1,498 | 1,157 | 12,887 |
| (%) | (26.0) | (25.5) | (23.8) | (24.9) | (25.6) | (24.7) | (25.7) |
| Q3, 15.45-21.55 | 7,412 | 1,257 | 179 | 866 | 1,342 | 1,171 | 12,227 |
| (%) | (24.9) | (22.9) | (24.7) | (23.4) | (23.0) | (25.0) | (24.4) |
| Q4, >38.52 | 7,715 | 1,370 | 185 | 874 | 1,260 | 1,104 | 12,508 |
| (%) | (26.0) | (24.9) | (25.6) | (23.6) | (21.6) | (23.6) | (24.9) |
| **Confirmation method of diagnosis** |  |  |  |  |  |  |  |
| Microscopic | 29,628 | 5,321 | 697 | 3,652 | 5,688 | 4,223 | 49,209 |
| (%) | (99.7) | (96.9) | (96.3) | (98.6) | (97.3) | (90.1) | (98.1) |
| Radiologic and clinic | 40 | 122 | 21 | 43 | 104 | 285 | 615 |
| (%) | (0.1) | (2.2) | (2.9) | (1.2) | (1.8) | (6.1) | (1.2) |
| Unknown/Other | 62 | 50 | <15* | <15* | 55 | 178 | 359 |
| (%) | (0.2) | (0.9) |  |  | (0.9) | (3.8) | (0.7) |
| **PSA, quartiles (ng/ml)** |  |  |  |  |  |  |  |
| <4.9 | 8,360 | 765 | 88 | 665 | 874 | 367 | 11,119 |
| (%) | (28.2) | (14.0) | (12.3) | (18.1) | (15.1) | (8.0) | (22.3) |
| 5.0-6.8 | 7,406 | 829 | 108 | 735 | 1,023 | 337 | 10,438 |
| (%) | (25.0) | (15.2) | (15.1) | (20.0) | (17.6) | (7.3) | (20.9) |
| 6.9-11.3 | 6,331 | 1,199 | 157 | 804 | 1,239 | 580 | 10,310 |
| (%) | (21.4) | (22.0) | (21.9) | (21.8) | (21.4) | (12.6) | (20.7) |
| 11.3+ | 4,081 | 1,487 | 216 | 887 | 1,494 | 2,331 | 10,496 |
| (%) | (13.8) | (27.3) | (30.1) | (24.1) | (25.8) | (50.6) | (21.1) |
| Unknown/Other | 3,433 | 1,168 | 148 | 590 | 1,170 | 992 | 7,501 |
| (%) | (11.6) | (21.4) | (20.6) | (16.0) | (20.2) | (21.5) | (15.0) |
| **Gleason score** |  |  |  |  |  |  |  |
| 5 | <15* | <15* | <15* | <15* | <15* | <15* | 15 |
| 6 | 264 | 37 | <15* | 21 | 26 | 19 | 374 |
| (%) | (0.9) | (0.7) |  | (0.6) | (0.5) | (0.4) | (0.8) |
| 7 | 219 | 29 | <15* | 22 | 28 | 32 | 335 |
| (%) | (0.7) | (0.5) |  | (0.6) | (0.5) | (0.7) | (0.7) |
| 8 | 36 | 15 | <15* | <15* | <15* | 20 | 94 |
| (%) | (0.1) | (0.3) |  |  |  | (0.4) | (0.2) |
| 9 | <15* | <15* | <15* | <15* | <15* | <15* | 65 |
| (%) |  |  |  |  |  |  | (0.1) |
| 10 | <15* | <15* | <15* | <15* | <15* | <15* | <15* |
| Unknown/Other | 29,069 | 5,358 | 701 | 3,624 | 5,728 | 4,493 | 48,973 |
| (%) | (98.2) | (98.4) | (97.8) | (98.5) | (98.8) | (97.5) | (98.2) |

Note: AJCC, 6<sup>th</sup> edition clinical staging of the American Joint Commission on Cancer; TNM6, 6<sup>th</sup> edition Tumor, node and metastasis staging manual of the American Joint Commission on Cancer; API, Asian Pacific Islanders; NH, Non-Hispanic; CVD, cardiovascular disease; PSA, Prostate specific antigen; \*, statistically suppressed; ¶ 95% confidence intervals in parenthesis; §, County attributes of Year 2000; Education attainment defined as percent of residents with less than high-school graduate in the county; Person in poverty defined as percent of residents with income below 200% of poverty in the county.

**Table 2.** Prediction accuracy for long-term 6-category causes of death among the patients with prostate cancer diagnosis in 2004 (follow up through Dec. 2016)

| Predicted classes | Alive,<br>n=14,746 | CVD,<br>n=2,689 | Infection,<br>n=371 | Non-Prostate<br>cancer,<br>n=1,873 | Other<br>cause,<br>n=2,897 | Prostate<br>cancer,<br>n=2,288 | Total,<br>n=24,864 |
| --- | --- | --- | --- | --- | --- | --- | --- |
| **Random forest model** |  |  |  |  |  |  |  |
| Alive, % | 87.70* | 52.73 | 52.29 | 67.49 | 55.82 | 39.9 | 73.75 |
| CVD, % | 3.79 | 15.88* | 15.90 | 10.04 | 15.08 | 8.92 | 7.54 |
| Infection, % | 0.21 | 0.67 | 0.27* | 0.32 | 0.69 | 0.31 | 0.33 |
| Non-Prostate cancer, % | 1.94 | 3.35 | 2.96 | 2.94* | 3.11 | 3.23 | 2.44 |
| Other cause, % | 3.82 | 17.44 | 16.44 | 10.62 | 15.05* | 10.01 | 7.87 |
| Prostate cancer, % | 2.54 | 9.93 | 12.13 | 8.60 | 10.25 | 37.63* | 8.06 |
| **Multinomial model** |  |  |  |  |  |  |  |
| Alive, % | 82.63* | 33.51 | 31.27 | 51.84 | 37.04 | 32.87 | 64.34 |
| NA, % | 17.37 | 66.49 | 68.73 | 48.16 | 62.96 | 67.13 | 35.66 |

Note: CVD, cardiovascular disease; NA, not available; \*, correct prediction.

### Predicting multi-category causes of death with random forests model

There were 17 variables with categorical encoding and 61 variables with one-hot encoding, and 240 candidate models in each tuning process. Our tuning processes showed that the prediction accuracy increased with the iteration number in either conventionally or one-hot encoded data (**Figure 2**), as shown before.[30] The mean prediction-accuracy for 6-category COD were 58.6% (95% CI, 58.2%-59.1%) in the RF models with conventional encoding and 59.1% (95% CI, 58.7%-59.4%) in those with one-hot encoding. The best accuracy was reached in the model of 3 variables and 800 iterations with conventional encoding (59.2%, 95% CI [58.6%-59.8%], Table 2 and **Figure 3**) and that of 1 variable and 700 iterations with one-hot encoding (59.6%, 95% CI [58.9%-60.2%], **STable 4** and **Figure 3**). The best RF model with one-hot encoding appeared to outperform that with conventional encoding, but no statistically significant difference was found. Alive was the COD that all RF models could predict with the best accuracy, while cancer pathological staging and age at diagnosis were top-important factors in the RF models (**Figure 3**).

**Figure 2.**
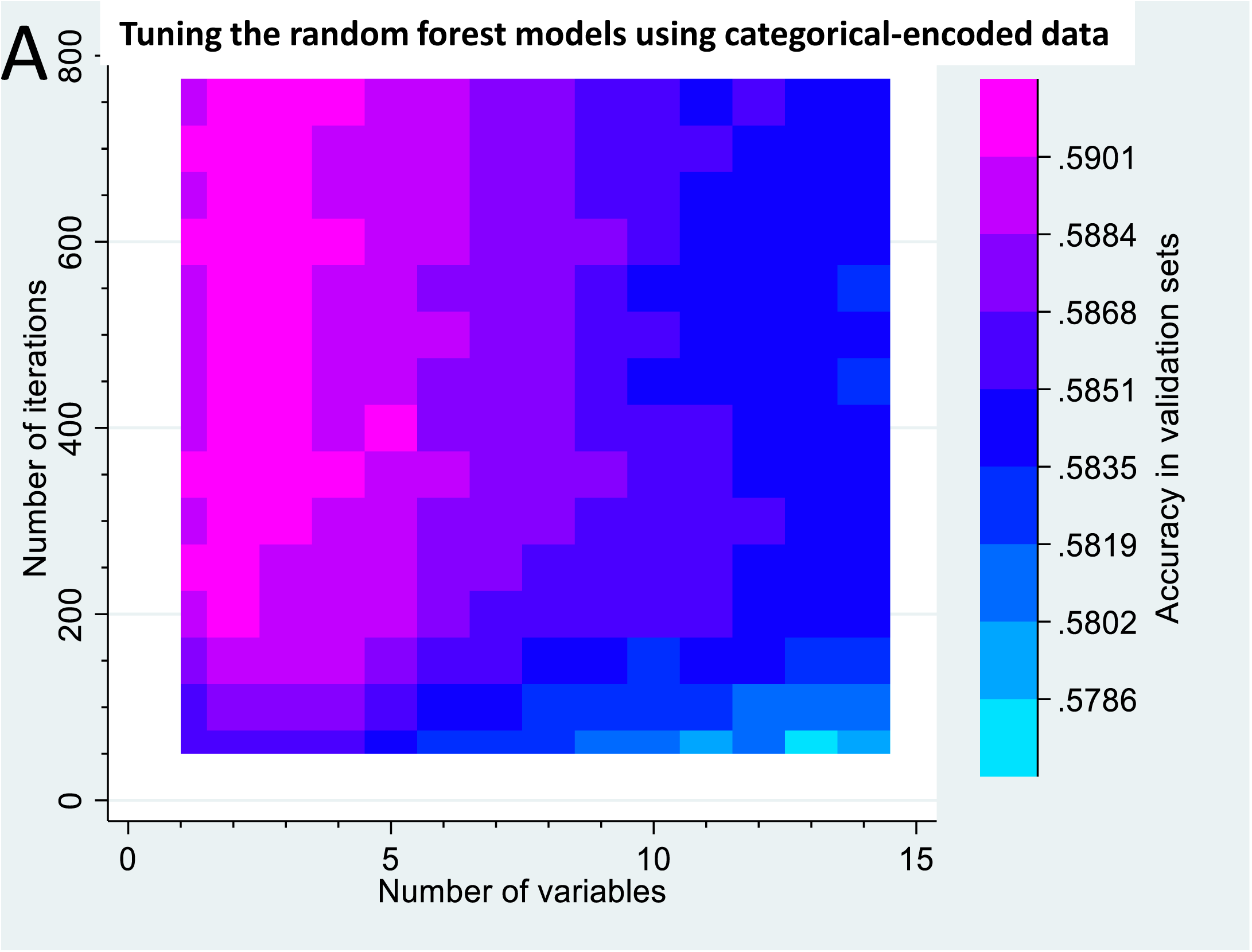
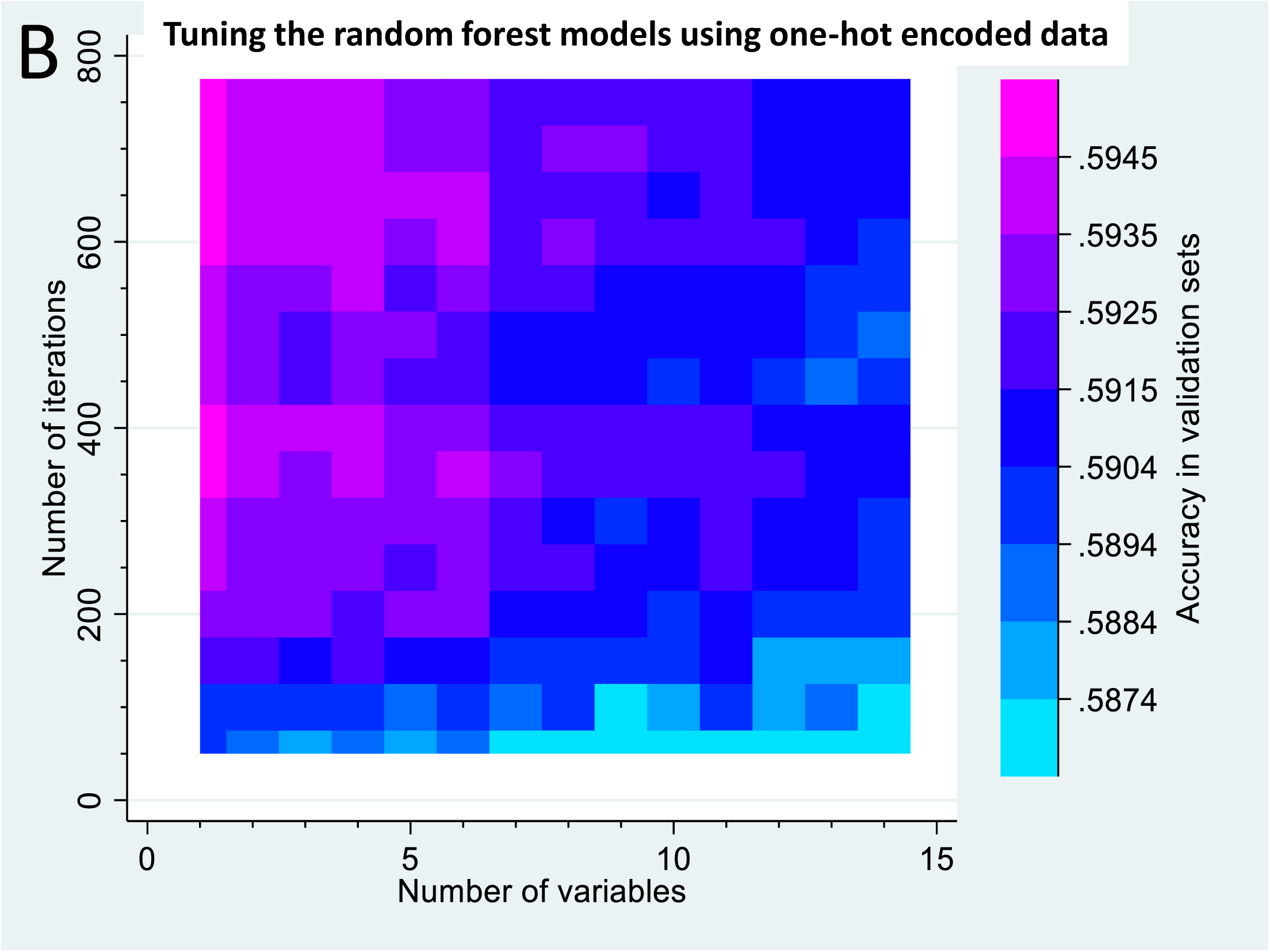
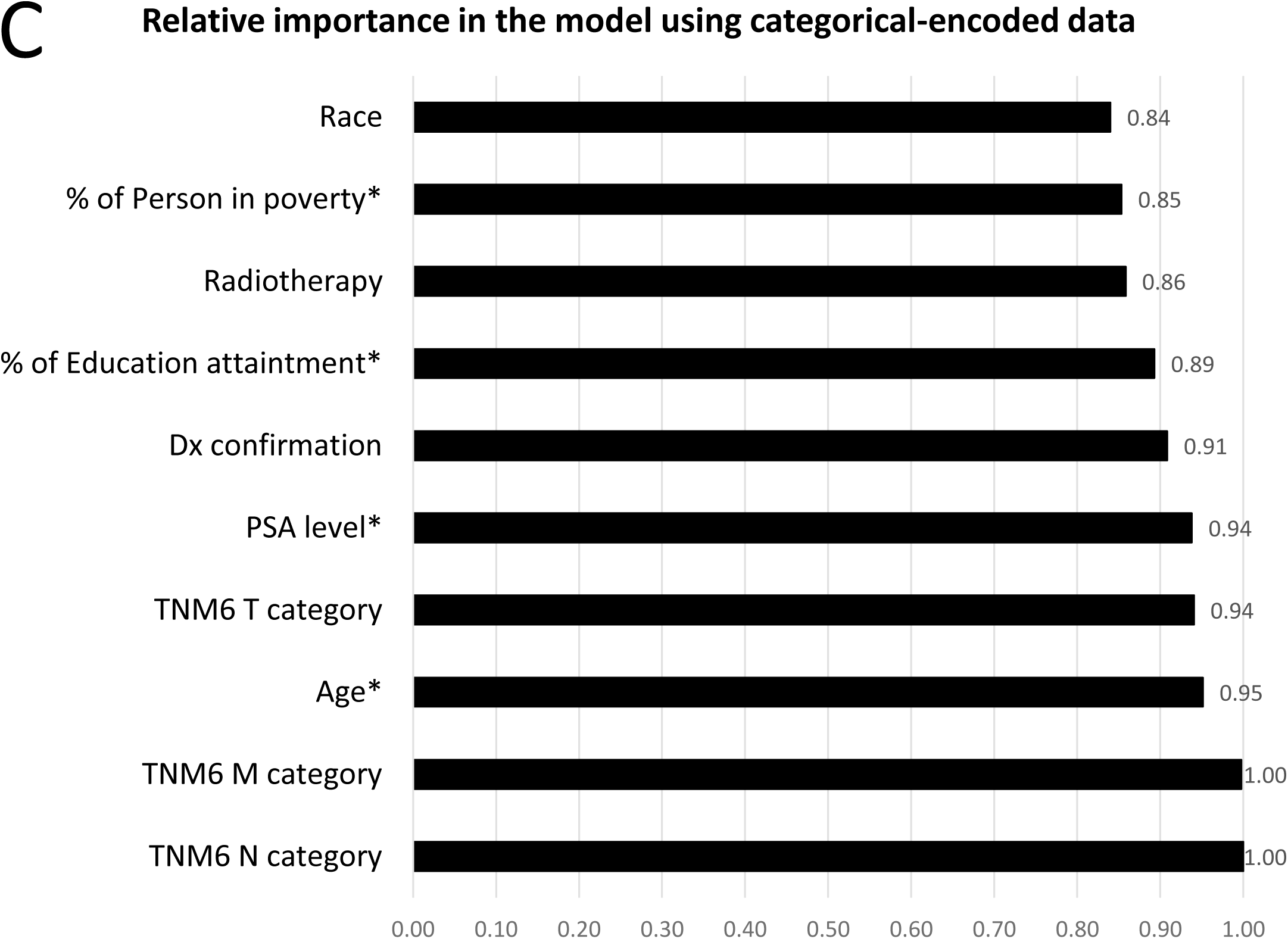
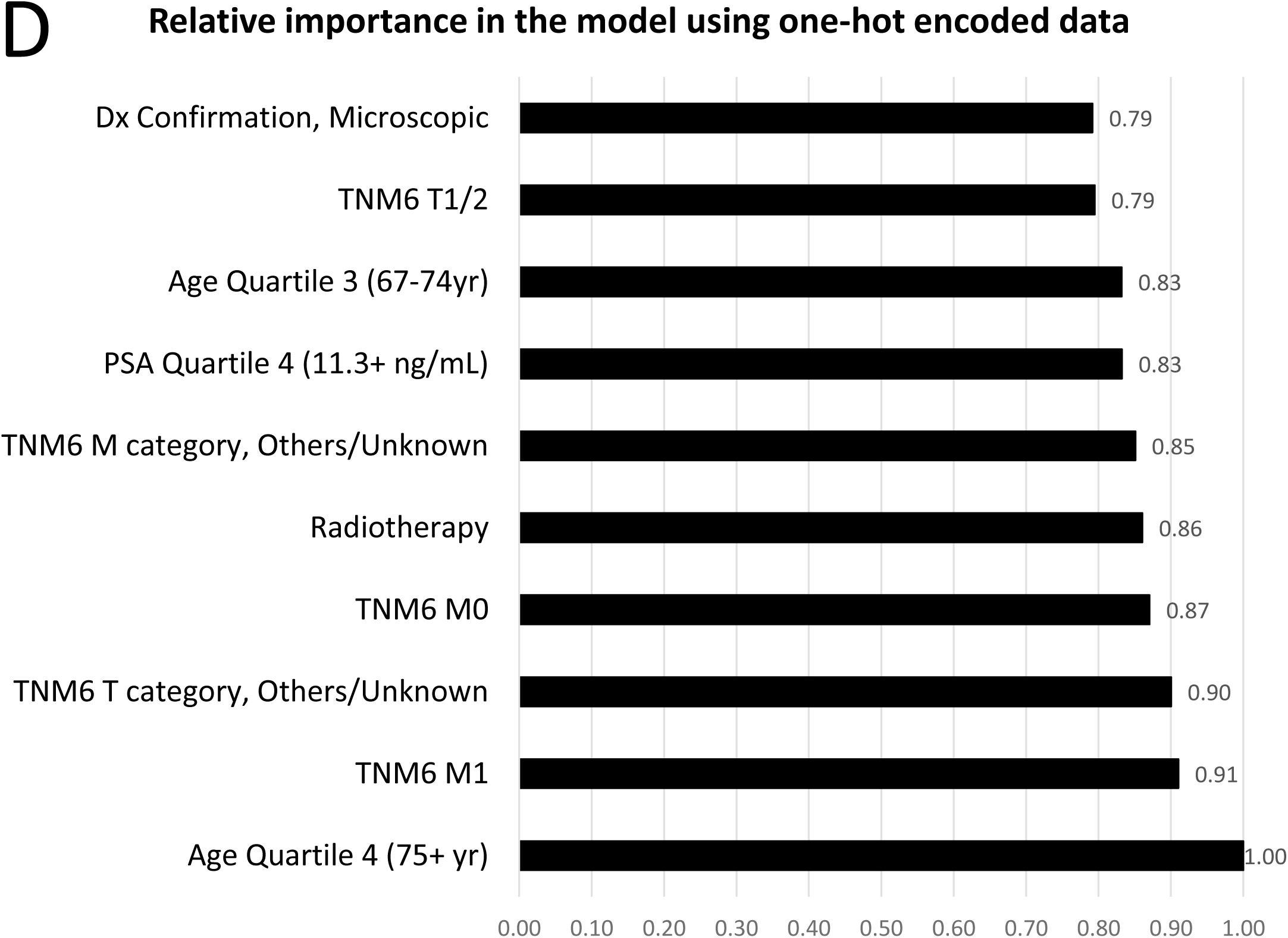
Characteristics of random forest models. During primary validation process, prediction accuracies of random forest models varied by the corresponding numbers of variable and iteration (Heatmap graphs: **A**. categorical data encoding; **B**. One-hot data encoding). The random forest models provided computed relative importance-values for all included variables (**C**. and **D**. Relative-importance values of the top 10 variables in the chosen random forest models using categorical data encoding and one-hot data encoding, respectively). Note: *, Continuous variables were converted to 4-category variables by their respective quartiles; Dx, diagnosis; PSA, Prostate specific antigen; Education attainment defined as percent of residents with less than high-school graduate in the county; Person in poverty defined as percent of residents with income below 200% of poverty in the county.

**Figure 3.**
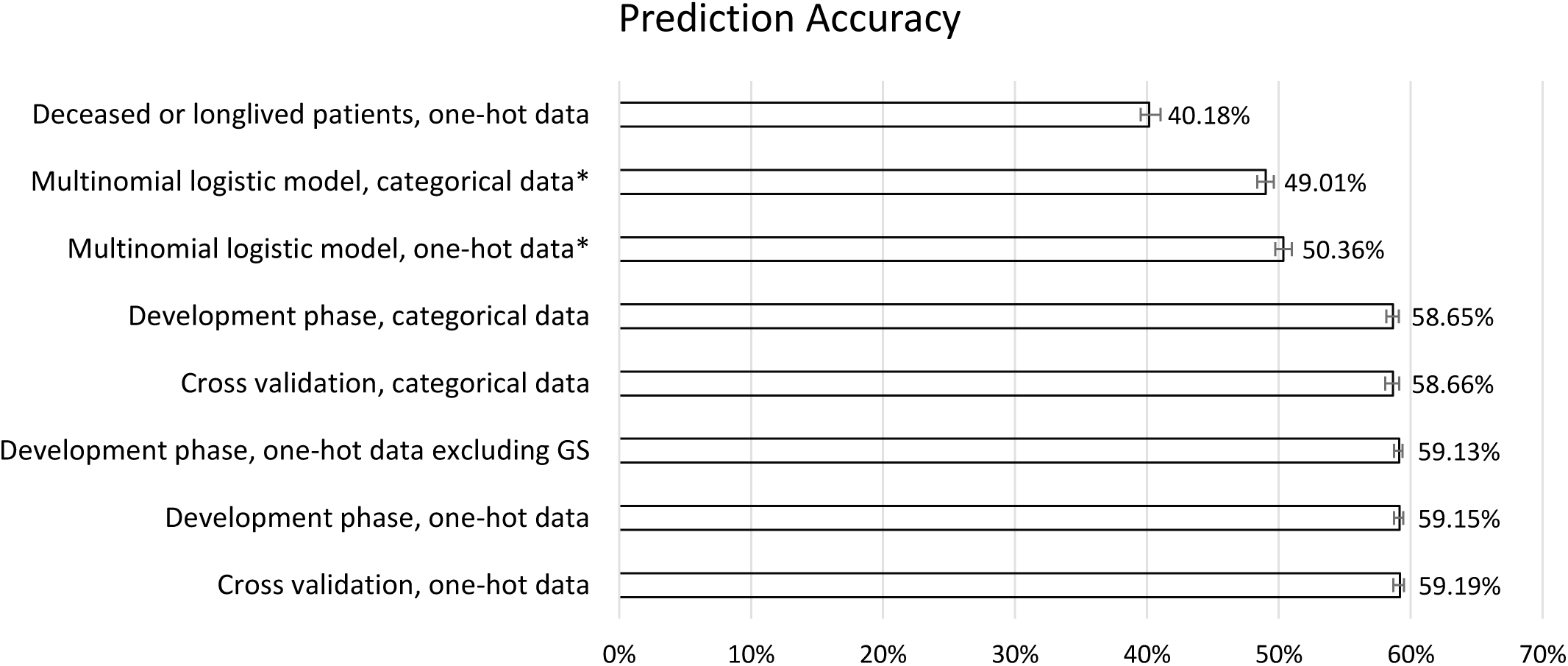
Summary of prediction accuracies by model and data type. In the tuning process and sensitivity analyses, we computed the validation accuracy of each random forest model by the numbers of variable and iteration (n=240), and chose the one with the best accuracy as the final model. The error bars show 95% confidence intervals of prediction accuracies in those models and data types during tuning process, except 3 models, whose 95% confidence intervals were calculated for the accuracy of a single model using binomial model (indicated by *). One-hot indicates one-hot encoding of the data; balanced set refers to the sensitivity analysis with training and testing sets that had balanced distribution of all variables.

The sensitivity analyses revealed that the prediction accuracies were statistically similar in the -training phase, and cross-validation phase, but statistically lower in the models in patients who died during follow-up or was alive for >150 months (**Figure 3**).

### Predicting multi-category causes of death with multinomial model

As the RF models, and the multinomial logistic regression models with one-hot encoding seemed to have better goodness of fit than with categorical encoding (Pseudo/adjusted R-square= 0.1707 versus 0.1416, Likelihood Ratio [Chi-square]= 10854.51 vs 9009.2, respectively). Because multinomial models used a ranking approach to determine the best-fit outcome, it is possible that more than one outcome (i.e. COD) had a probability >0.5. However, the predicted COD in multinomial model was only unique in being alive among the 6-category COD and all other categories were of < 0.5 probability (**Table 2** and **STable 4**). The mean prediction-accuracy was 50.4% (95% CI, 49.7%-51.0%) in the multinomial models, and lower than RF models, except the RF model on the patients who died during followup or was alive for >150 months (**Figure 3**). Age at diagnosis, AJCC6 staging, confirmation method of diagnosis, surgery and PSA level were associated with all 6-category COD in multinomial model, while other factors were only linked to some of the 6-category COD (**STable 5**).

## Discussion

In this study, we investigated the multilevel prediction problem of prostate patients’ COD using a carefully constructed and rigorously tuned RF model. In the patients with prostate cancer diagnosed in 2004, 59.4% were alive at the end of 12-year follow-up, while the top-3 CODs were CVD, prostate cancer and non-prostate cancer. We predicted 6-category COD among these patients with a mean accuracy of 59.1% (95% CI, 58.7%-59.4%) in the tuned RF model with one-hot encoding, and 50.4% (95% CI, 49.7%-51.0%) in the multinomial logit model, suggesting RF models outperformed multinomial model. Tumor characteristics, PSA level, diagnosis confirmation-method, and radiotherapy status were the top-ranked variables in RF model, but only age, surgery, diagnosis confirmation-method, PSA level and AJCC6 stages as the factors were linked to all of the COD (versus alive) in multinomial models.

The proportions of various COD in our study are similar to those in previous reports.[4] Given the increasing proportion of deaths from COD other than prostate cancer, it is critical to accurately predict or identify the factors linked to these COD among prostate cancer patients. Several studies have attempted to predict cancer-specific or all-cause deaths in prostate cancer patients using clinical pathological and genomic/genetic factors.[15, 33-36] However, few studies to our knowledge predict the causes of death in multiple categories. Multinomial logistic regression is suitable for analyzing categorical/multi-category outcomes.[31, 32] In this study, multinomial logistic regression seems only able to predict the alive status of the 6-category COD if a unique COD successfully identified. In the meantime, a tuned RF model outperformed multinomial logistic regression in predicting 6-category COD by 17.2% higher prediction accuracy (8.7% absolute accuracy-difference). This finding supports that RF’s accuracy is similar to or better than support vector machines, artificial neural network and logistic regression in predicting various clinical outcomes,[9-11, 37] but contrasts to that its accuracy is inferior to that of logistic regression.[38] It is plausible, but needs additional validation, that RF could also be highly useful in predicting multi-category COD or outcomes of other diseases. Despite the slightly better accuracy linked to data with one-hot encoding than standard encoding, we found no statistical differences between the two methods. This finding is inconsistent with previous reports,[28, 29] and needs further validation. We also noticed that the minimal depths of trees in our best-fit RF models were usually 1 to 3. Those observations may help develop and improve machine learning models for predicting multi-category COD in cancer or other patients.

Some of this study’s strengths are noteworthy. First, this population-based study provides early evidence on the frequencies of various COD among the prostate cancer patients who were followed up for 12 years. Second, we tuned RF models for predicting 6-category COD in prostate cancer patients, while existing RF models on prostate cancer only predicted binary cancer-specific death,[15, 33] all-cause death,[33, 39] or cancer recurrence.[40] Compared with binary death-outcomes, multiple-category COD are more informative, but more difficult to predict. This is supported by the low success rate of multinomial models in predicting unique COD. Third, the tuned RF models in this study outperformed multinomial models in predicting 6-category COD. Indeed, the multinomial model was only able to predict alive as a unique COD, and missed other COD. Fourth, we characterized RF models and identified the model with best accuracy, while few of the prior works tuned their models.[15, 33, 40] Fifth, we are able to able to achieve a promising prediction accuracy given the large sample-size of this prostate cancer dataset and the cross-validation.[41] Some of prior studies on prostate cancer survivals using machine learning/RF model had either large sample size[15] or cross validation,[42-44] but few combined both. Small sample size was indeed reported as the most common limitation of machine learning studies on cancer prognosis and prediction.[41] Finally, we identified several factors linked to long-term 6-category COD in prostate cancer patients, including age, PSA level and tumor characteristics, as shown by both RF and multinomial models.

This study has the following limitations. The prediction accuracy for 6-category COD in this study is not yet as good as prediction for binary outcomes, such as all-cause deaths.[33] Moreover, despite some shared linked-factors, RF models did not completely agree with multinomial models on the factors linked to 6-category COD. However, RF and other machine learning models are known for their limitations in identifying associated factors.[45] In addition, an outside validation dataset might be needed, but unavailable, largely due to the lack of registry-data. SEER18 is the largest population cancer dataset in the North America.[16] Thus, it is very challenging to obtain another population dataset of similar size for validation. However, we prospectively used the cross-validation approach to validate our findings, as recommended.[41, 45] Finally, Gleason scores were available in a very small proportion of the patients, but might otherwise improve prediction accuracy.[46]

## Conclusions

In this population-based study, CVD, prostate cancer and non-prostate cancer were the most common long-term COD among prostate cancer patients. RF and multinomial models could predict 6-category COD among these patients with acceptable prediction accuracy, which needs improvement. Those models enable clinicians to gain more granular prognostic information on prostate cancer patients, and target at relevant COD to improve survival. We also show that a tuned RF model outperforms multinomial models by 8.7% (absolute difference), or 15,195 person-case for the cases diagnosed in 2019 alone in the U.S. Additional works are needed to better predict multiple-category COD of other cancers.

## Statement

### Availability of data and material (Mandatory)

All data are available through SEER after signing an agreement.

### Authors’ contributions

JW, FZ, AJS and LZ designed the study, JW, FD and LZ extracted and analyzed the data, JW and LZ wrote the first draft of the manuscript and all authors edited the manuscript. The final manuscript was approved by all authors except FZ, who unfortunately passed away on Oct. 1, 2018.

## Key Points

- A tuned random forest model could reach an accuracy of 59.1% in predicting long-term 6-category cause of death among U.S. prostate cancer patients.
- Random forest model outperforms conventional-statistical/multinomial models by an absolute percent difference of 8.7%, but its accuracies did not differ by data coding (conventional versus one-hot encoding).
- Tumor characteristics, prostate-specific antigen level, and diagnosis confirmation-method appear important for predicting 13-year 6-category cause of death in both random forest and multinomial models.

