## Supplementary figures and images for "Predicting long-term multicategory cause of death in patients with prostate cancer: random forest versus multinomial model"

### Supp fig 1

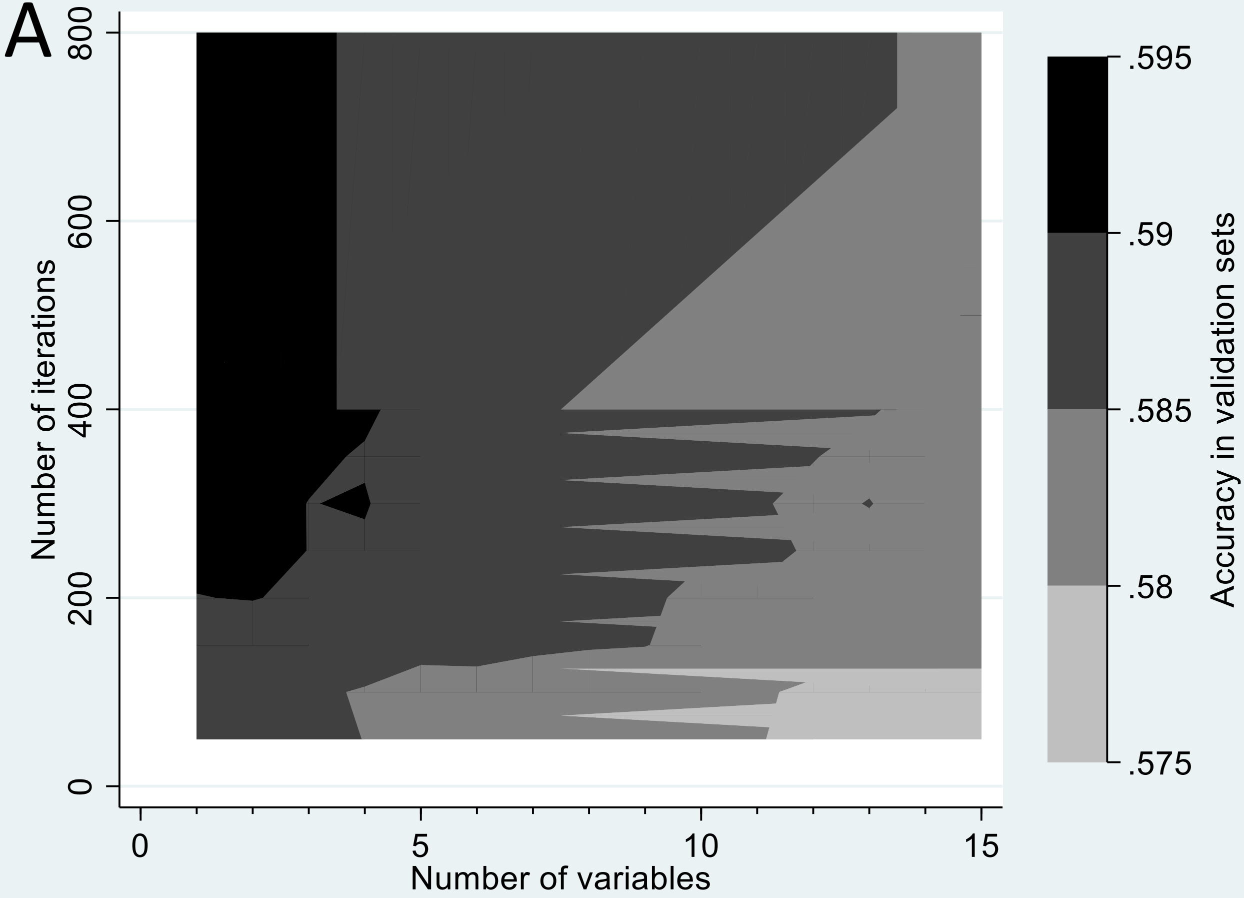

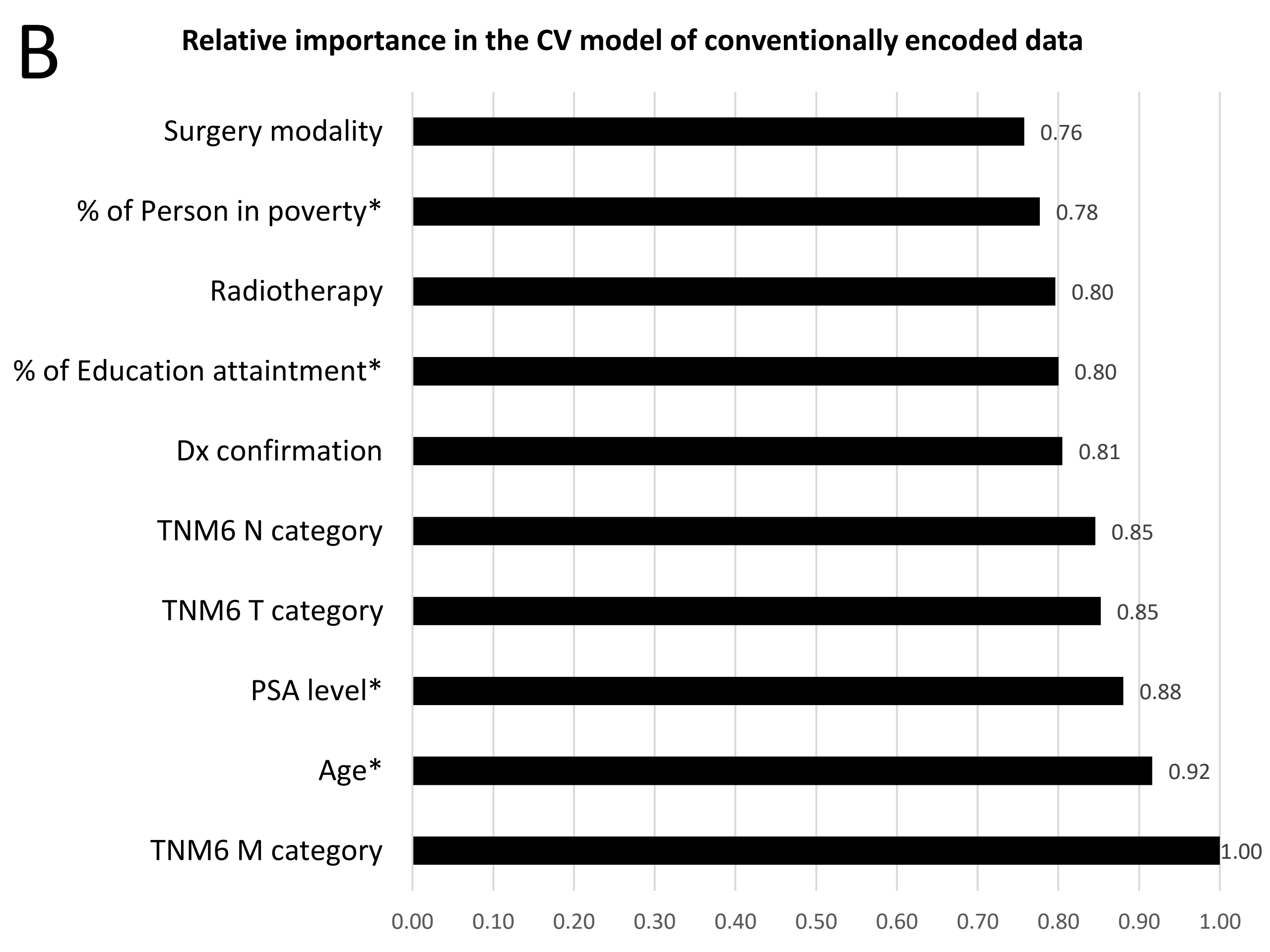
